# Pervasive and dynamic gut dysbiosis in wild bumble bees

**DOI:** 10.64898/2026.02.05.704103

**Authors:** Kristal M. Watrous, McKenna J. Larson, Bahareh Sorouri, Ozichi Ikegbu, Annika S. Nelson, Jonathan B.U. Koch, Tobin J. Hammer

## Abstract

Stressors can shift the microbiome into an altered, “dysbiotic” state that reduces host fitness. While well-studied in humans and laboratory models, the prevalence, predictability, and drivers of dysbiosis in wildlife remain unclear. We addressed these questions by monitoring gut microbiomes of wild bumble bees in Southern California, focusing on *Bombus vosnesenskii*, a major pollinator in western North America. More than a third of all *B. vosnesenskii* bees exhibited dysbiosis, when defined as a >50% replacement of host-specialized core bacteria by environmental bacteria. This replacement covaried with increased alpha and beta diversity, an enrichment of oxygen-tolerant taxa, and pathogen infection—all common hallmarks of dysbiosis in other hosts. Other co-occurring *Bombus*, including two at-risk species, also exhibited dysbiosis. In *B. vosnesenskii*, dysbiosis was not correlated with certain stressors, such as heat or resource limitation, although other, unmeasured stressors cannot be ruled out. We next examined how dysbiosis varied over two years of sampling. In the first year, dysbiosis emerged only late in the season, when bumble bee colonies normally reproduce and then senesce. Two years later, following an intervening year with historic rainfall and elevated resources, dysbiosis was entrenched throughout the season. These findings show that dysbiosis is both pervasive and highly dynamic in wild bumble bees. The dynamics coincide with host life cycle transitions and environmental change, but the underlying causality remains uncertain. Given that dysbiosis may harm host health, we argue that long-term microbiome monitoring should be considered both for bumble bees and for other wildlife of conservation concern.

## Introduction

Perturbations that alter the microbiome beyond the bounds of its normal dynamics can reduce host health and fitness. Such destabilized, harmful microbiome states are often referred to as dysbiosis^1–4^. Research on humans and experimental models has shown that dysbiosis can arise when an extrinsic stressor disrupts mechanisms of host control, including immunity or physicochemical conditions, that act to maintain microbiome stability^5,6^. However, distinguishing healthy from unhealthy forms of microbiome change, or natural from anthropogenic, is not straightforward^7–9^. For example, some of the conventional hallmarks of dysbiosis—altered microbial diversity, increased interindividual variation, loss of “core” taxa, enrichment of pathogenic taxa, and inflammation^10–12^—can arise in healthy humans during normal life cycle transitions, such as reproduction or aging^13–16^.

Characterizing dysbiosis is particularly difficult in wildlife, because the environment is inherently variable, and information on the traits and life histories of individual hosts is usually limited. Even in closely monitored populations of wild vertebrates, microbiome surveys often find extensive interindividual and temporal microbiome variation without obvious phenotypic correlates^17–20^. In cases where the microbiome correlates with disease or other markers of host health^17,21–23^, it is difficult to identify whether the microbiome affects health, or whether the causality is reversed or due to an independent factor (e.g., diet). Indeed, the conceptual underpinnings of dysbiosis have been criticized for these and other reasons^9,24–26^. But determining the causes and consequences of microbiome change in wildlife remains an important challenge, regardless of the terminology used. Phenomena such as coral bleaching, amphibian chytridiomycosis, or white-nose syndrome in bats illustrate how shifts in the microbiome can disrupt host populations and ecosystem function^27–29^.

Bumble bees (*Bombus* spp.) are a unique model for studying the ecology of dysbiosis. The adult stage harbors a relatively simple (in terms of membership) and clearly definable core gut microbiome, dominated by a few bacterial symbiont lineages within the genera *Snodgrassella*, *Gilliamella*, *Schmidhempelia*, *Lactobacillus*, *Bombilactobacillus*, *Bifidobacterium*, and *Bombiscardovia*^30,31^. These lineages are defined as “core” not based on an arbitrary prevalence threshold, as is often the case in other systems^32^, but by their host-restricted nature^31,33^. Bumble bee-associated core bacterial strains cannot colonize the honey bee gut effectively^34,35^ and likely cannot grow in the environment at all^33^. They have close relatives in honey bees and stingless bees, suggestive of an ancient symbiosis dating back to the common ancestor of the corbiculate Apinae^30,36,37^. Further, gnotobiotic experiments with *B. terrestris* and *B. impatiens* have shown that the core microbiome provides colonization resistance against eukaryotic and bacterial pathogens^38–40^ and can also modulate behavior^41^ and phytochemical metabolism^42^. When *B. terrestris* and *B. impatiens* are reared in captivity, the core bacteria comprise nearly all of the gut microbiome of individual worker bees^43,44^.

Paradoxically, these ancient and important symbionts are sometimes missing from wild bumble bees. Field surveys consistently uncover bees in which the expected core gut bacteria have been partially or entirely replaced by non-host-specific, environmental bacteria^31^. Such bees, from a wide range of *Bombus* species, have been documented in Asia, Australia/New Zealand, Europe, North America, and South America, and in protected natural areas as well as urban and agricultural environments^40,45–53^. The environmental bacteria-dominated gut microbiome state has variously been labeled as dysbiosis, an alternate enterotype, or microbiome disruption^31,45–47,49,53,54^. Whether this state indeed fits under the general concept of dysbiosis has been unclear, because its characteristics have not been explicitly compared with common definitions or examples in other hosts. For simplicity, and because it is justified by our results, we use the term dysbiosis below.

Dysbiosis may pose a threat to bumble bees and the organisms that depend on them. Bumble bees are ecologically and agriculturally important pollinators^55^, but populations of several species have declined or have gone extinct^56,59,61,63,79,80^. The potential role of dysbiosis in these declines is unknown^31^. While no obvious phenotypes or pathologies have been reported, three lines of evidence suggest that dysbiosis is harmful: loss of benefits from the core microbiome (noted above), pathogenicity of at least some of the environmental bacteria^40^, and associations between dysbiosis and eukaryotic gut pathogens (*Crithidia* or *Nosema* [syn. *Vairimorpha*]) in wild populations^45,46,49^. Superficially at least, bumble bee dysbiosis resembles coral bleaching, a phenomenon where host-specific microbial symbionts are disrupted, impairing host health. Bleaching has catastrophic impacts not only for individual corals but for entire reef ecosystems, motivating interventions to increase resilience of the symbiosis^57^.

One hypothesis is that dysbiosis is ubiquitous in wild bumble bees because it is driven by ubiquitous environmental stressors. For example, extreme heat, suboptimal food resources, and pesticides can disrupt bee immunity or gut microbiomes^58,60,62,64^. There is ample precedent for this kind of mechanism. Extrinsic stressors are well-known to induce gut dysbiosis and other microbiome perturbations in humans, experimental models, and some wild animals^18,65–71^. Further, as the major stressors affecting bees are either a direct result of human activity or are exacerbated by climate change^72^, support for this hypothesis would suggest that bumble bee dysbiosis is an anthropogenic phenomenon.

Alternatively, dysbiosis may be ubiquitous because it is an intrinsic feature of bumble bee life history. Most bumble bee species are eusocial and have an annual life cycle, which begins with colony founding by a solitary queen. The colony then undergoes a growth phase, during which multiple cohorts of worker bees are produced. Subsequently, sexuals (males and new queens) are produced, disperse from the colony, and mate. Finally, the nest is abandoned and all of the bees die, excepting the new queens, which undergo diapause (i.e., overwinter) alone in underground hibernacula^55,73,74^. Studies with captive-reared *B. terrestris* in Europe have found that environmental bacteria replace core bacteria in the gut of worker bees as colonies age, suggesting a link between dysbiosis and the colony life cycle^44,75^. However, the predictability and field-relevance of this pattern are unclear, because very few studies have tracked gut microbiome dynamics over the colony cycle in wild populations. Furthermore, environmental stressors may increase over the same seasonal time scale as the colony cycle^76,77^. Overall, it remains unclear whether dysbiosis is intrinsic to bumble bee biology, anthropogenic, or perhaps—like coral bleaching^78^—an intrinsic stress response that is increasingly triggered by anthropogenic environmental change.

To provide insight into the dynamics and drivers of dysbiosis in wild bumble bees, we monitored gut microbiomes of a *Bombus vosnesenskii* population in Southern California, USA over the colony cycle in two years. *B. vosnesenskii* is a highly abundant, stable species in the western USA^79^, making it useful for consistent monitoring. We measured microbial composition and abundance across 155 worker and male *B. vosnesenskii* bees and used metagenomics to characterize the identities and traits of dominant bacterial taxa. To place microbiome change in the context of the bees’ life history and environment, we also collected data on phenology, weather, resource availability, and several host traits. Three co-occurring *Bombus* species were sampled in the second year to assess the broader extent of dysbiosis. These included *B. melanopygus*, *B. sonorus*, and *B. californicus*; the latter two, belonging to the *B. pensylvanicus* and *B. fervidus* species complexes, respectively, are listed as “Vulnerable” by the IUCN^80^. Our results demonstrate that dysbiosis may be widespread in wild bumble bees, with dynamics linked to both the host life cycle and to environmental change.

## Results & Discussion

### Dysbiosis in wild Bombus vosnesenskii

Gut microbiomes of our focal bumble bee (*Bombus vosnesenskii*) population, sampled across the colony cycle in both 2022 and 2024, were highly variable in composition and host specificity (Fig. 1A). Some individuals exclusively harbored the core, bumble bee-restricted gut bacterial symbionts known from other *Bombus* species^30,31,81^. Metagenome-assembled genomes (MAGs) of core taxa from *B. vosnesenskii* have the highest sequence identity to isolates from the guts of other *Bombus* (Table S1), corroborating their host-specificity. These individuals’ gut microbiomes resemble those of honey bees (*Apis* spp.) in terms of being fairly consistent among individuals and dominated by core bacteria^82,83^. However, 37% of individuals harbored microbiomes with >50% environmental bacteria (Fig. 1A). All of these genera occur in plants, solitary bees, honey bees, and/or other insects^84–89^, suggesting flower-mediated bacterial transmission into bumble bees. The relative abundance of environmental bacteria captures much of the overall variation in microbiome composition (Fig. S1; linear models vs. NMDS Axis 1; 2022, N = 60, R^2^ = 0.71, p < 0.001; 2024, N = 95, R^2^ = 0.69, p < 0.001). Hence, we use it as a simplified measure of microbiome composition for many of the following analyses.

**Figure 1.**
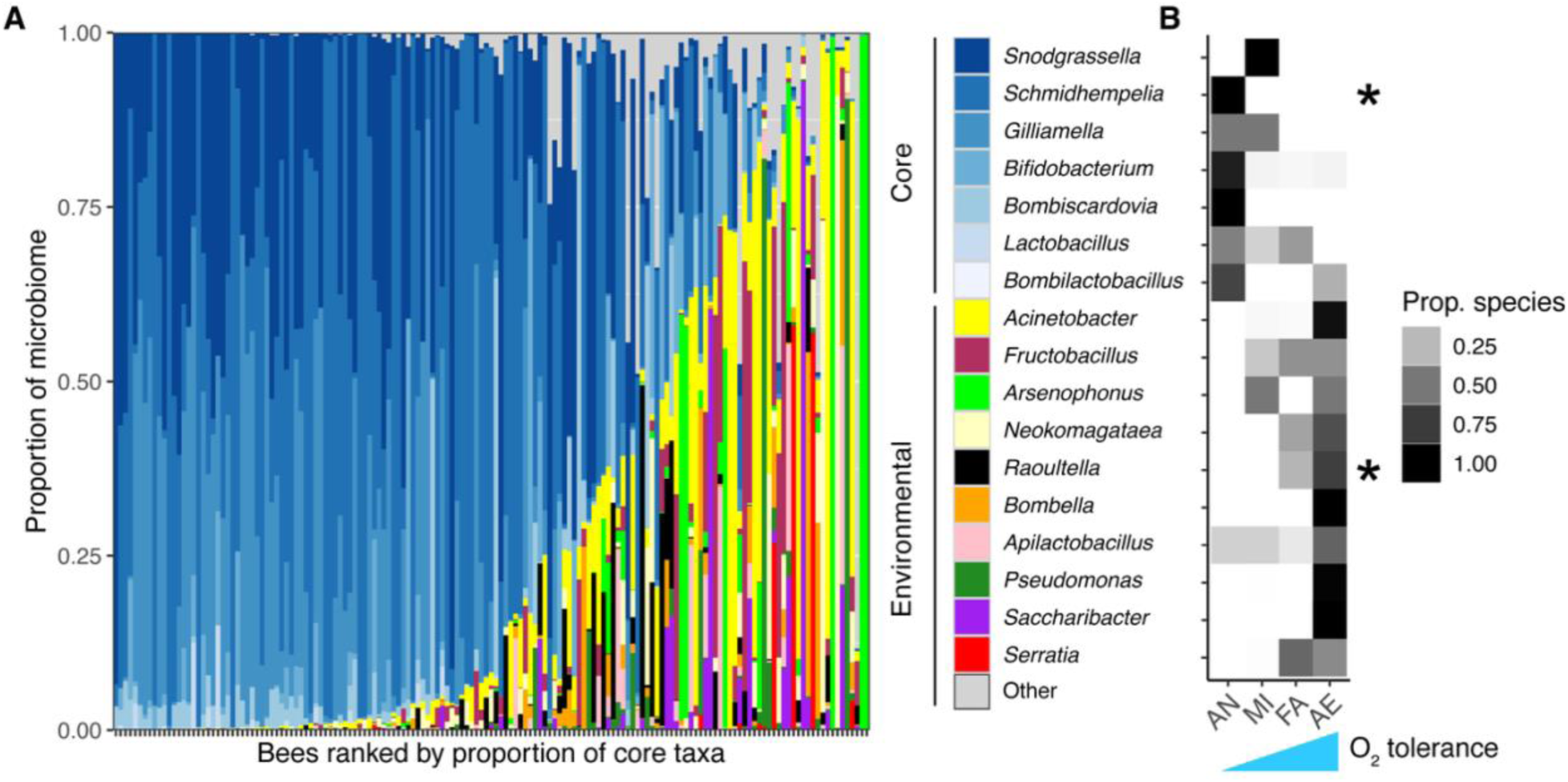
Prevalence and oxygen tolerance of dominant gut bacterial genera in wild bumble bees (*B. vosnesenskii*). A) Gut microbiomes vary widely in the proportion of core (bumble bee-specialized) bacteria versus environmental bacteria, as measured by 16S rRNA gene sequencing. Each column is an individual bee sampled from the same population in 2022 (N = 50 workers, 10 males) or 2024 (N = 71 workers, 24 males). Patterns broken down by sex and year are shown in Fig. 3A and Fig. 4A. “Other” (gray) represents all environmental bacterial genera with <1% mean relative abundance across all samples. B) Most species belonging to core bacterial genera cannot grow under atmospheric oxygen levels—i.e. are anaerobic, AN, or microaerophilic, MI—while most environmental bacteria either tolerate (facultative aerobe, FA) or require it (obligate aerobe, AE). Rows correspond to the genera shown in Fig. 1A, in the same order. The shading (white to black) represents, for a given bacterial genus, the proportion of species in the BacDive database classified to a particular oxygen-tolerance category. Asterisks indicate values that were manually added based on other sources because the genera are absent from BacDive (see Methods).

The ten most common environmental bacteria (Fig. 1A) are phylogenetically diverse (representing seven families in four orders), but one trait that unifies them, and sets them apart from the core bacteria, is oxygen tolerance. Oxygen tolerance varied between core and environmental bacterial genera (Fisher’s exact test, N = 17, p < 0.001), with core genera comprising mostly anaerobic or microaerophilic species, and environmental genera comprising mostly facultatively anaerobic or aerobic species (Fig. 1B). In honey bees, the core bacteria are necessary to maintain an anoxic gut lumen^90^. This effect may be driven by bacterial fermentation byproducts fueling host epithelial cell respiration^91^, or by the microaerophile *Snodgrassella*, which forms a biofilm on the ileum wall in both honey bees and bumble bees^31,90^. Possibly, if the core bacteria are displaced (or never acquired), oxygenation of the gut inhibits their subsequent (re)establishment, giving a competitive edge to oxygen-tolerant environmental bacteria.

The balance of core versus environmental bacteria is strongly associated with disease. *B. vosnesenskii* workers enriched in environmental bacteria have a much higher probability of being infected with the trypanosomatid gut pathogen *Crithidia* (binomial GLM: N = 122, odds ratio = 14.30 [95% CI = 4.37, 53.53], z = 4.16, p < 0.001; Fig. 2A). One worker harbored *C. expoeki*, while another 23 (out of 122 total) harbored *C. bombi*, as determined by blastn searches; none of the 34 males were infected with either *Crithidia* species. *C. bombi* has been shown to strongly reduce bumble bee (*B. terrestris*) survival and reproduction, although only under stressful conditions^92,93^. This result is congruent with field data from various *Bombus* species wherein depleted levels of core bacteria are correlated with infection by *C. bombi* and other eukaryotic pathogens^45,46,49^. While the causality remains unknown for wild *Bombus* populations, laboratory experiments with *B. terrestris* and *B. impatiens* demonstrate the potential for bidirectional feedbacks. These experiments have shown that disrupting colonization by the core microbiome strongly increases susceptibility to *C. bombi*^38,39^. Further, *C. bombi* may itself disrupt the core gut microbiome^94^ (but see ^95^), an effect that could be mediated by host immune system activation^96,97^.

**Figure 2.**
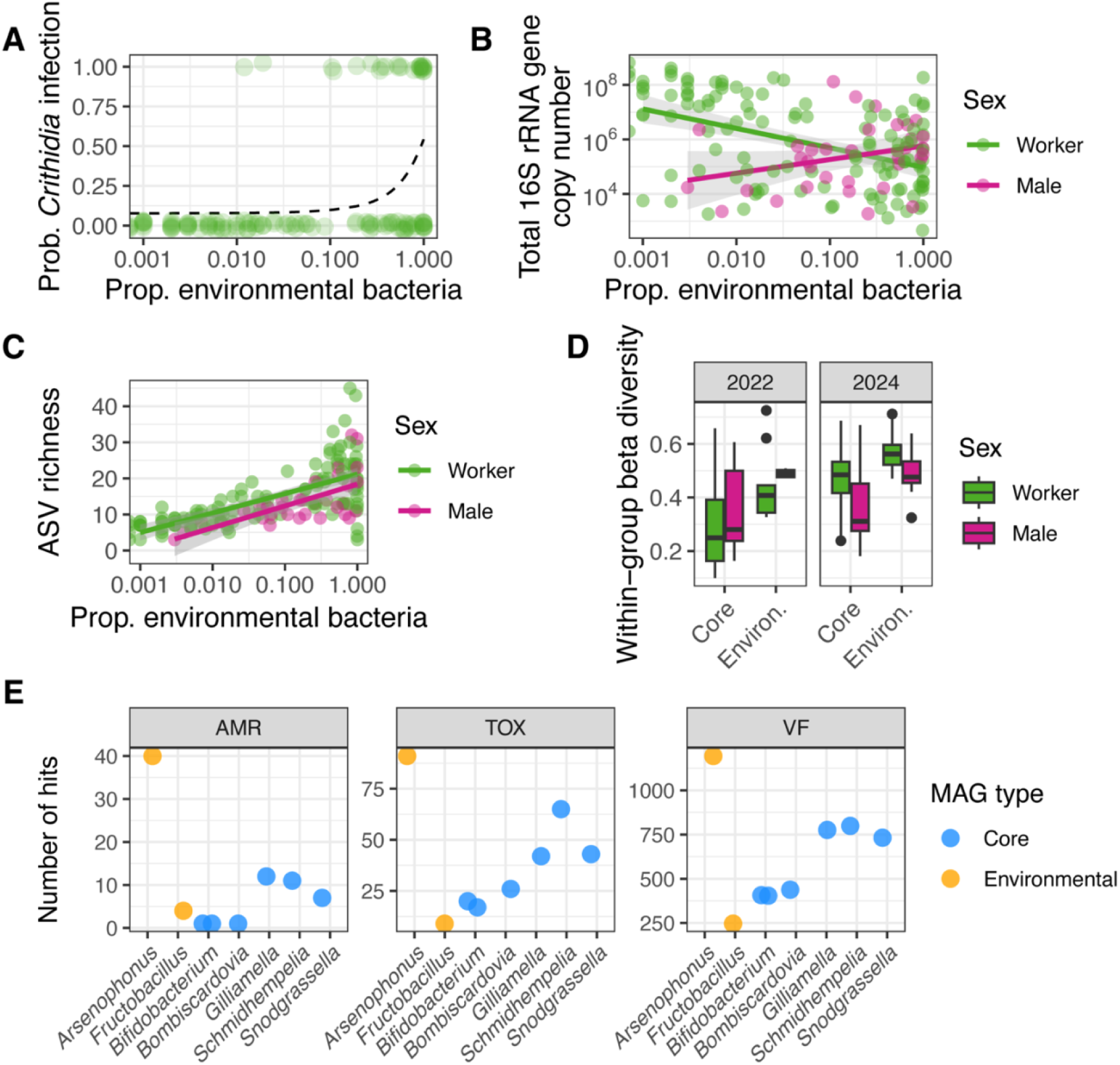
Dominance of environmental bacteria in the *B. vosnesenskii* gut microbiome presents features that, alongside those shown in Fig. 1, are characteristic of dysbiosis. The x axes in panels A-C are log-transformed to facilitate visualization. A) The proportion of environmental bacteria in worker bee microbiomes predicts infection by the trypanosomatid pathogen *Crithidia*. The dashed line shows a logistic regression model fitted to the data. Points are vertically jittered to reduce overplotting. *Crithidia* was not detected in males. B) Worker bees with higher proportions of environmental bacteria have less overall bacterial biomass, as measured using qPCR with universal 16S rRNA gene primers. Lines show linear models fitted to the data and 95% confidence intervals in gray. C) Dominance of environmental bacteria is associated with elevated alpha diversity (richness of bacterial amplicon sequence variants) in workers and males. D) Dominance of environmental bacteria is associated with elevated beta diversity (interindividual variation in microbiome composition). Samples are binned by majority-core or majority-environmental sequences. Sample sizes are as in the Fig. 1A caption. Horizontal lines indicate medians, box limits indicate first and third quartiles, and whiskers extend to the most extreme values within 1.5X the inter-quartile range. E) The genome of *Arsenophonus*, a common environmental bacterium in *B. vosnesenskii*, is enriched in antimicrobial resistance (AMR), toxins (TOX), and virulence factors (VF). Shown are the number of hits identified by PathoFact for eight metagenome-assembled genomes (MAGs) from *B. vosnesenskii* (Table 1).

Some of the environmental bacteria common in *B. vosnesenskii* are themselves likely capable of opportunistic pathogenicity. *Serratia* has been experimentally demonstrated to be pathogenic to bumble bees (*B. impatiens*), as has *Pseudomonas* in honey bees, and *Arsenophonus* has been linked to honey bee colony collapse disorder and mortality^40,98–100^. The *Arsenophonus* MAG from *B. vosnesenskii* has high average nucleotide identity (ANI) to an isolate from honey bee hemolymph (Table S1), suggesting this bacterium can escape the bee gut and cause septicemia. Further, it is enriched in three classes of putative pathogenicity-related factors (Fig. 2E). It should be noted, however, that many molecules labeled as virulence factors or toxins are common to beneficial or neutral symbionts as well as pathogens^101,102^. The *B. vosnesenskii*-associated *Fructobacillus* MAG does not exhibit high numbers of pathogenicity-related factors (Fig. 2E), but it has high ANI to an isolate from a honey bee larva with foulbrood disease (Table S1), suggesting the ability to colonize the bee gut upon perturbation.

The amplicon sequence data shown in (Fig. 1A) are proportional (relative abundance), an attribute that can obscure microbiome dynamics. To assess the relationship between gut bacterial biomass (absolute abundance) and microbiome composition, we used quantitative PCR with universal primers to estimate total 16S rRNA gene copy numbers. Bacterial biomass (16S copy numbers) and the proportion of environmental versus core bacterial sequences are strongly associated across bees, but to a degree that depends on sex (linear model [LM], N = 155, bacterial composition * sex interaction: p = 0.0057; Fig. 2B). Worker bees with more environmental bacteria have lower total 16S copy numbers (LM: N = 121, R^2^ = 0.12, t = −4.15, p < 0.001), while in males, there is no significant relationship (LM: N = 34, t = 1.26, p = 0.22). We interpret this as showing that, in workers, lost or missing core bacterial biomass is not fully offset by environmental bacterial biomass. As environmental bacteria are unlikely to be adapted to the bumble bee gut^31,33,103^, their growth may be suppressed to a greater degree (as compared with core bacteria) by host immunity or other factors.

Next, we examined how the balance of core versus environmental bacteria corresponds to ecological diversity metrics. Alpha diversity (the number of unique sequence variants within a given bee gut) was strongly predicted by the proportion of environmental bacteria in both workers and males (LM: N = 155, R^2^ = 0.34, t = 8.94, p < 0.001; Fig. 2C). To compare beta diversity (inter-individual variability in community composition), we grouped individuals based on whether the majority of gut bacterial sequences belonged to core or environmental bacteria, and on year of collection. Beta diversity varied among these groups (ANOVA on distances to group centroids; N = 155, F = 15.15, p < 0.001; Fig. 2D). Post-hoc pairwise tests show that beta diversity was higher in environmental-dominated workers versus core-dominated workers in both years (adjusted p < 0.05); the same comparison was not significant for males, potentially due to low sample sizes. These ecological diversity patterns show that shifts between core and environmental-dominated microbiomes are not from one simple, stable state to another. Rather, the absence of core bacteria seems to destabilize the microbiome, allowing colonization by a comparatively diverse and erratic set of taxa.

The characteristics of the environmental-dominated microbiome of wild *B. vosnesenskii* bumble bees form a syndrome with clear parallels to gut dysbiosis as described in humans and laboratory models. Commonalities include the loss of core bacteria, acquisition of oxygen-tolerant and potentially pathogenic bacteria, association with disease (*C. bombi*), altered alpha diversity, and increased compositional variability. Further, this syndrome fits with various conceptual frameworks for dysbiosis, such as those emphasizing loss of core taxa, destabilization, or disruption of host control^3,6,10–12,104,105^. Thus, bumble bees provide one of the few well-characterized examples of dysbiosis in nature, and could serve as a model for recognizing and studying this phenomenon in other wildlife.

### No evident role of diet or foraging effort in dysbiosis

In honey bees and bumble bees, resource limitation and composition affect the gut microbiome^40,60,94,106^ and conversely, the gut microbiome can affect foraging and feeding behavior^90,107^. Therefore, we sought to determine whether diet and dysbiosis are associated in *B. vosnesenskii*. We first assessed resource limitation, using the number of pollen grains in the gut as a measure of recent consumption^108,109^. As expected from work on *B. impatiens*^109^, *B. vosnesenskii* males had less pollen in their gut than workers (LM: N = 155, R^2^ = 0.16, t = 5.59, p < 0.001; Fig. S2A). However, there was no significant relationship between ingested pollen and dysbiosis in either workers (LM: N = 121, t = 1.70, p = 0.092) or males (N = 34, t = 1.44, p = 0.16) (Fig. S2A). We next assessed resource composition. Plant taxa vary in traits such as nectar secondary metabolites and pollen morphology, and laboratory experiments have shown that these traits can affect pathogen establishment in the gut of *B. terrestris* and *B. impatiens*^40,42,109^. However, we found that bees with core-dominated microbiomes foraged on a similar distribution of plant families as bees with environmental-dominated microbiomes (Fig. S2B). A random forest classifier trained to predict plant family from dysbiosis and sex performed poorly (N = 153, OOB error rate: 50.98%). Further, dysbiosis was less useful for prediction than sex (mean decrease in accuracy of 33% when permuting sex, versus 15% when permuting dysbiosis). Overall, these results suggest that bumble bee dysbiosis is not strongly associated with diet.

Bumble bee workers may spend hours per day flying to collect food while carrying loads up to three quarters of their body mass^110,111^, an activity that imposes metabolic costs, generates oxidative stress, and can reduce immune responses^112–114^. We examined whether foraging effort underlies *B. vosnesenskii* dysbiosis, using the degree of wing wear as a proxy^115^. Wing wear varied extensively among individuals, as expected from studies of bumble bee foraging behavior^110,116^, but was not associated with dysbiosis (ordinal regression: N = 155, z = −0.45, p = 0.65) nor with sex (z = 0.58, p = 0.56) (Fig. S2C). It should be noted that some bumble bee workers do not leave the nest^110,116^; these would not be represented among our sampled bees, which were all collected while foraging. However, the results are consistent with a previous study using captive-reared but free-foraging *B. terrestris* colonies, in which the gut microbiome did not differ between foragers and non-foragers^44^.

### Seasonal dysbiosis covaries with the life cycle

If bumble bee dysbiosis is not caused by extrinsic stressors, an alternative hypothesis—with some precedent from human gut microbiome studies^13–16^—is that it is an intrinsic feature of host life cycle transitions. To assess support for this hypothesis, we analyzed temporal gut microbiome dynamics in *B. vosnesenskii* from early spring until late summer, starting in 2022. Sampling began when workers were first observed and continued biweekly until they were no longer observed. In 2022, gut microbiome composition varied over time (db-RDA: N = 60, F = 7.62, p < 0.001; Fig. S1A;), with worker bees transitioning from core-dominated to environmental-dominated (i.e., dysbiotic) states toward the end of the season (beta regression: N = 60, z = 3.90, p < 0.001; Fig. 3A). Not all of the core bacterial taxa exhibited the same dynamics. For example, *Snodgrassella* and *Bifidobacterium* persisted through the end of the season, while all of the others virtually disappeared (Fig. S3). As observed previously in wild *B. impatiens*^40^, *Schmidhempelia* dominated the microbiome of early-season *B. vosnesenskii* workers (Fig. S3), likely because it primarily colonizes young bees (at least in *B. impatiens*) ^43^.

**Figure 3.**
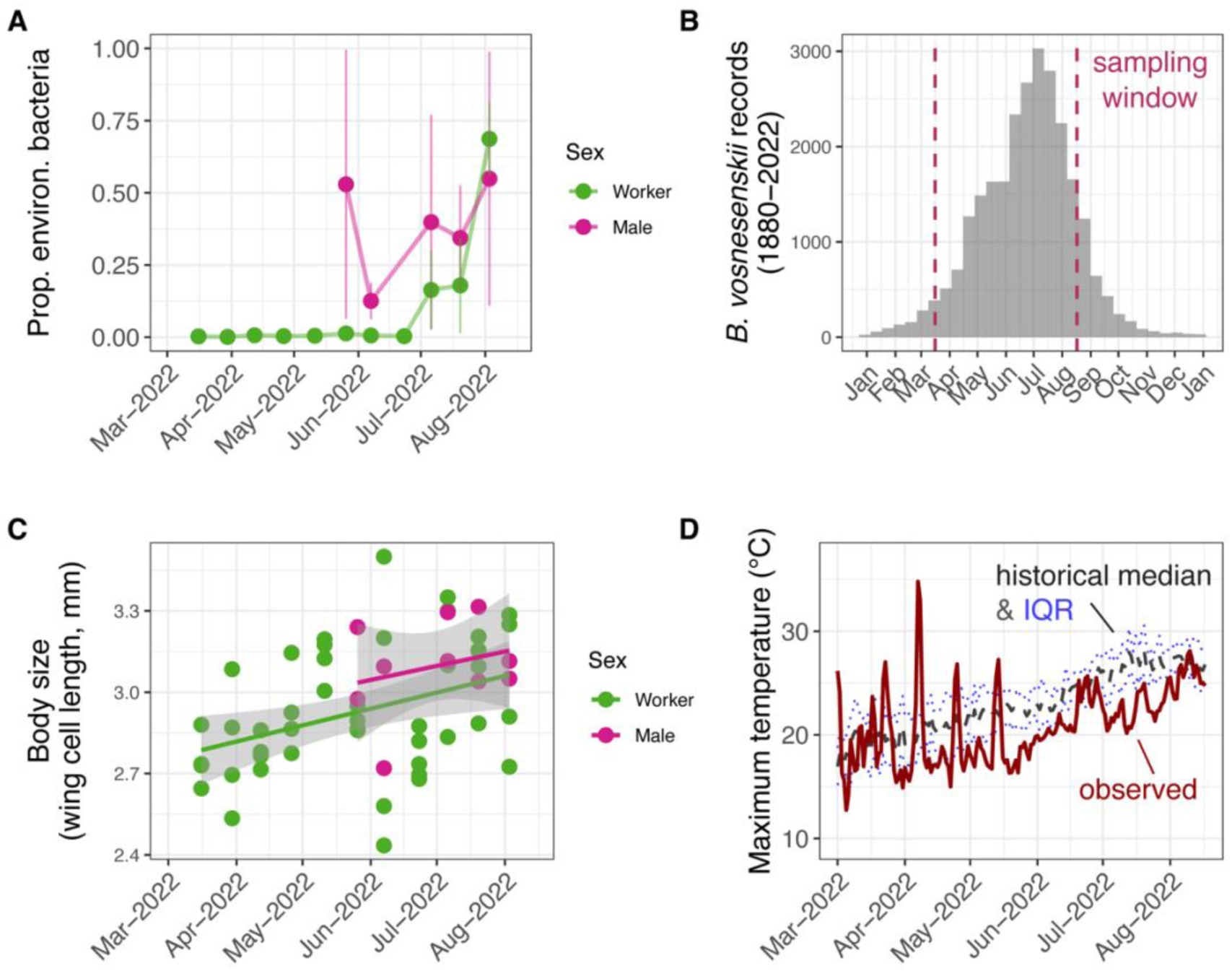
In 2022, *B. vosnesenskii* dysbiosis coincided with the expected end of the bumble bee colony life cycle. Tick marks on the x axis correspond to the first day of each month. A) Dysbiosis (quantified as a high proportion of environmental bacteria) only emerged in worker bees toward the end of the season. Males are produced later in the colony cycle and did not show temporal variation. Points represent means of replicate bees sampled on the same day and vertical lines represent the SEM (N = 50 workers, 10 males). B) The sampling period (dashed lines) spanned the typical phenology of *B. vosnesenskii*, wherein populations annually transition from growth to reproduction to senescence. Bars show the number of *B. vosnesenskii* specimens (any sex or caste) collected in California in a given time of year, from 1880-2022. C) Worker and male size increased over the sampling period, as would be expected to occur over the colony cycle. The averaged length of the left/right forewing marginal cells was used as a proxy for body size. D) Dysbiosis does not appear to be driven by heat stress. Daily maximum temperatures at the field site in coastal Southern California were obtained from the PRISM Daily Spatial Climate Dataset. Solid line: observed temperatures in 2022; dashed line: 30-year historical median temperature; dotted lines: interquartile range of historical temperatures.

It is plausible that a pattern of seasonal microbiome change at the population level could emerge from sampling different bee colonies over time, where each colony has a fixed microbiome. In order to differentiate this pattern from within-colony dynamics, we used genotyping to assign bees to putative colonies (wherein workers and males are expected to be siblings). Only a fraction of bees were inferred to be siblings of other bees in the sample set, making it difficult to quantitatively evaluate the two scenarios. However, for the seven colonies represented by multiple bees, we do find qualitative support for within-colony transitions from core to environmental gut microbiomes (i.e., transitions between siblings sampled over time) (Fig. S4A).

Males maintained compositionally distinct gut microbiomes from workers (db-RDA: N = 60, F = 6.02, p < 0.001; Fig. S1A), with a higher relative abundance of *Schmidhempelia* (LM: N = 60, t = 3.09, p = 0.003; Fig. S3) and a stronger tendency towards dysbiosis (beta regression: N = 60, z = 2.24, p = 0.025; Fig. 3A). The core bacteria as a whole appear to colonize males less effectively (Fig. 2B). Based on laboratory experiments with *B. impatiens*, this may reduce males’ resistance to invasion by environmental bacteria^40^. For bumble bees generally, males differ from workers in several traits that could influence the gut microbiome: they are produced, on average, later in the colony cycle; they spend much more time outside of the nest; they consume less pollen (Fig. S2A); and they may have weaker immunity^55,109,117^.

The late-season emergence of dysbiosis in 2022 (Fig. 3A) coincides with the end of the annual colony life cycle. In California, *B. vosnesenskii* population size generally peaks around July and then declines, based on historical collection records (Fig. 3B). Conventional bumble bee life history and studies of individual colonies of various species including *B. vosnesenskii*^116,118^ indicate that this population-level trend is driven by colony growth, followed by reproduction and senescence. Further evidence that our sampling window tracked the local *B. vosnesenskii* colony cycle comes from the observation that worker body size increased seasonally (LM: N = 50, R^2^ = 0.20, t = 3.64, p < 0.001; Fig. 3C). As individual *Bombus* colonies grow, worker body size tends to increase^119,120^.

Heat exposure does not explain the seasonal pattern of dysbiosis. Although daily maximum temperatures did increase over the season, they mostly stayed within or below the range of historical climate (Fig. 3D). Further, they did not approach typical thermal maxima of either bumble bees (CTmax^121^) or core gut bacteria^122^, as measured from various *Bombus* species. All of the anomalously warm days occurred in the spring (Fig. 3D), well before dysbiosis became prevalent in the *B. vosnesenskii* population.

The seasonal microbiome dynamic in wild *B. vosnesenskii* in 2022 is qualitatively similar to prior studies and involves many of the same environmental bacteria. In Europe, core bacteria were replaced by *Fructobacillus* or Enterobacteriaceae in older colonies of captive-reared *B. terrestris* placed outdoors^44,75^. In the northeastern United States, wild populations of multiple *Bombus* species showed repeated late-season shifts from core bacteria to *Fructobacillus*, Enterobacteriaceae, *Pseudomonas*, *Saccharibacter*, and *Acinetobacter*^49^. In an era of accelerating environmental change, it is difficult to rule out the possibility that a given microbiome dynamic is anthropogenic, particularly in the absence of historical evidence. It should be noted that our study region experienced severe to exceptional drought during the summers of 2013, 2014, 2015, 2016, 2018, 2021, and 2022 (U.S. Drought Monitor, droughtmonitor.unl.edu). Nevertheless, the recurrence of seasonal dysbiosis across *Bombus* species and regions suggests that the pattern we observed in 2022 may be intrinsic to bumble bee life history.

Why would dysbiosis arise toward the end of the colony cycle? Senescence of individual bees appears not to be a major factor, given our wing wear results (Fig. S2C) and prior studies of *B. terrestris* and *B. impatiens*^43,44^. Although speculative, one hypothesis is that, once colonies complete reproduction and begin to senesce, workers stop investing time and energy in cooperative behaviors^73^, such as hygiene or thermoregulation, that may normally reduce microbial contamination in the nest. A further consideration is that the core microbes cannot be lost at the end of every colony cycle—nor persist in host lineages over evolutionary time scales^30,37^—without a way to reclaim dominance in the gut. This microbiome “reset” may normally be mediated by the new queens produced at the end of the colony cycle, which transmit microbes from one generation to the next^31,123^. In *B. lantschouensis*, core gut bacteria were found to be depleted in young, pre-diapause queens, but abundant in queens that had completed diapause and founded new nests^124^.

### Environmental variability and entrenched dysbiosis

If late-season shifts from core to environmental bacteria are a typical feature of the annual life history of bumble bees, one would expect them to recur every year in the same population. To test this, we repeated biweekly sampling of the *B. vosnesenskii* population in 2024. The same suite of core and environmental bacteria that were dominant among gut microbiomes in 2022 reappeared in 2024 (Fig. 1A), and overall community composition again varied over the season in both workers (db-RDA: N = 71, F = 2.54, p = 0.009) and males (N = 24, F = 3.30, p = 0.004) (Fig. S1B). Further, worker body size increased over the season as in 2022 (LM: N = 71, R^2^ = 0.11, t = 3.04, p = 0.003), suggesting that the 2024 sampling period also tracked the colony life cycle^119,120^. However, the dynamics of dysbiosis completely changed. In 2024, environmental bacteria were already present in early-season worker bees and persisted until the end of the sampling period in September (beta regression: N = 71, z = 1.18, p = 0.26; Fig. 4A). Strangely, as in workers in 2022 (Fig. 3A), males in 2024 seasonally shifted from core to environmental bacteria (beta regression: N = 24, z = 4.56, p < 0.001; Fig. 4A). We again genotyped bee specimens to partition microbiome variability into separate colonies based on sibship. Unlike in 2022 (Fig. S4A), colonies in 2024 did not consistently switch from core to environmental bacteria over time. Rather, they displayed multiple patterns, including shifts from environmental to core bacteria as the season progressed (Fig. S4B).

**Figure 4.**
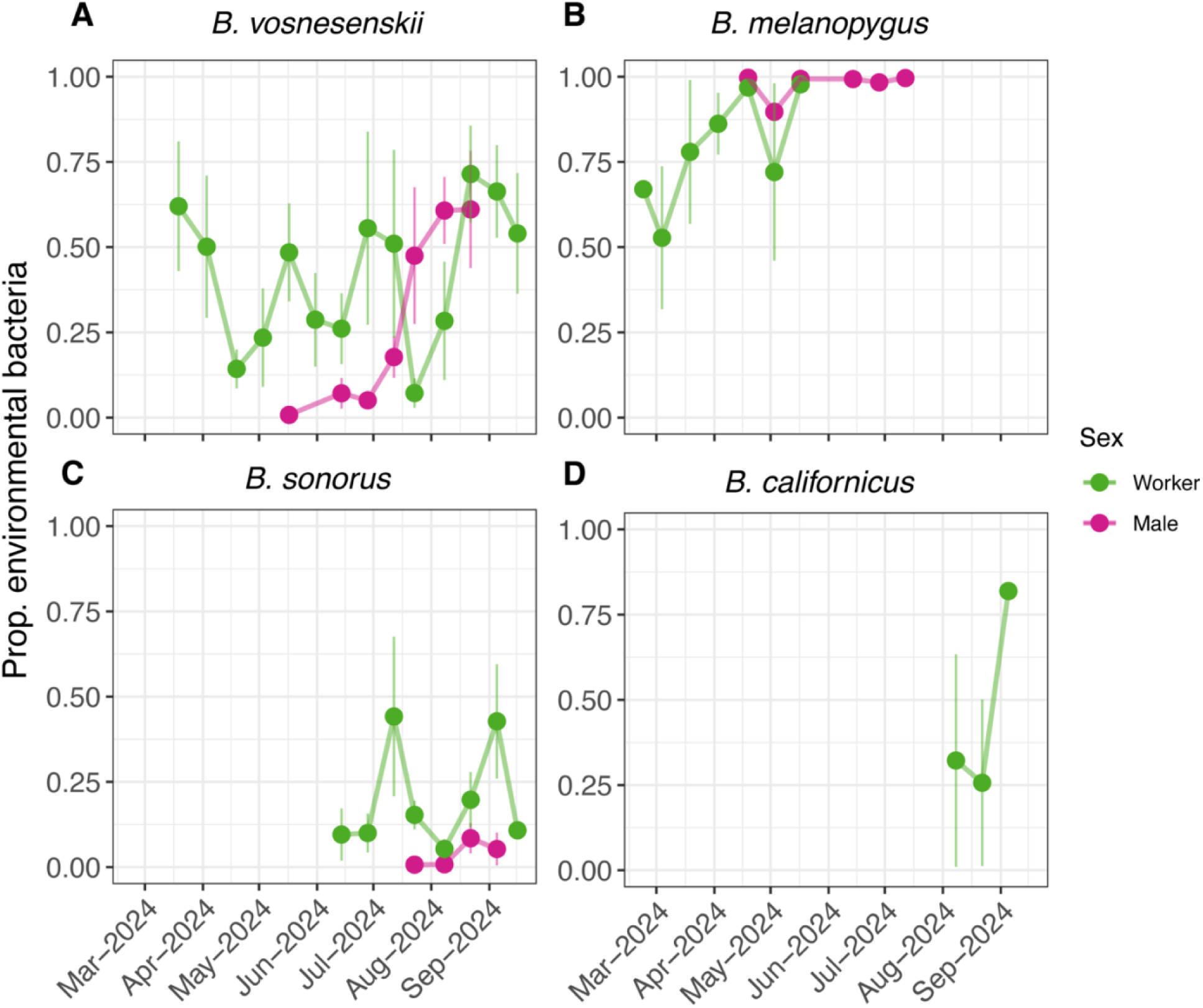
In 2024, dysbiosis was pervasive in *B. vosnesenskii* (A) and three co-occurring *Bombus* species (B-D). Species vary in the timing of collection because of differences in their phenology. Tick marks on the x axis correspond to the first day of each month. Points represent means of replicate bees sampled on the same day and lines represent the SEM. Sample sizes: A) N = 71 workers, 24 males; B) N = 27 workers, 12 males; C) N = 30 workers, 9 males; D) N = 8 workers.

Other, co-occurring *Bombus* species also exhibited dysbiosis, but at different frequencies. Since sample sizes for these species are much lower than for *B. vosnesenskii*, we describe the patterns qualitatively. Most *B. melanopygus* microbiomes were dysbiotic: 81% of workers (N = 27) and 100% of males (N = 12) harbored > 50% environmental bacteria (Fig. 4B). Dysbiosis was less prevalent in *B. sonorus*, where only 13% of workers (N = 30) and none of the males (N = 9) harbored > 50% environmental bacteria (Fig. 4C). Three out of eight *B. californicus* workers were dysbiotic (Fig. 4D). Although longer time series data are needed, the tentative observation that each *Bombus* species has a distinct pattern suggests that species-specific traits (e.g., phenology or nesting ecology) may modulate dysbiosis dynamics.

Eukaryotic gut pathogens were also widespread beyond *B. vosnesenskii*. 50% of *B. melanopygus* individuals (workers and males combined) harbored *Crithidia* (N = 42). For *B. sonorus*, 10.2% of individuals harbored *Crithidia*, and 12.8% harbored *Nosema* (syn. *Vairimorpha*) (N = 39), a microsporidian pathogen that, like *Crithidia*, reduces bumble bee (*B. terrestris*) fitness^125^. 37.5% of *B. californicus* harbored *Nosema* (N = 8). As in *B. vosnesenskii*, the relative abundance of environmental bacteria strongly predicted the presence of parasites (*Crithidia* and/or *Nosema*) in *B. sonorus* (binomial GLM: N = 39, odds ratio = 406.79 [95% CI = 11.44, 91594.36], z = 2.75, p = 0.005). It did not in *B. melanopygus* (binomial GLM: N = 39, odds ratio = 0.40 [95% CI = 0.021, 4.77], z = −0.70, p = 0.47), likely because so many individuals had microbiomes with very high levels of environmental bacteria (Fig. 4B). These results provide support for a consistent link between gut bacterial dysbiosis and pathogens in bumble bees, including in the at-risk species *B. sonorus*^80,126^.

If bumble bee dysbiosis were caused by stressors, it might be expected to become more common after periods with stressful conditions. Instead, we found that the “outbreak” of dysbiosis in 2024 (Fig. 4) followed an unusually favorable year for bumble bees, in terms of weather and resources. During the winter of 2022/2023, California received a series of atmospheric rivers that brought heavy precipitation^127^. At our field site, winter rainfall increased substantially from 2021/2022 to 2022/2023 and remained high in 2023/2024 (Fig. 5A). Strong rainfall led to elevated and sustained plant productivity, as inferred from the satellite imagery-derived normalized difference vegetation index (NDVI), a measure of vegetative greenness (Fig. 5B). In this region, more total plant productivity can be expected to result in more floral resources, and in turn, more bumble bees^118,128,129^.

**Figure 5.**
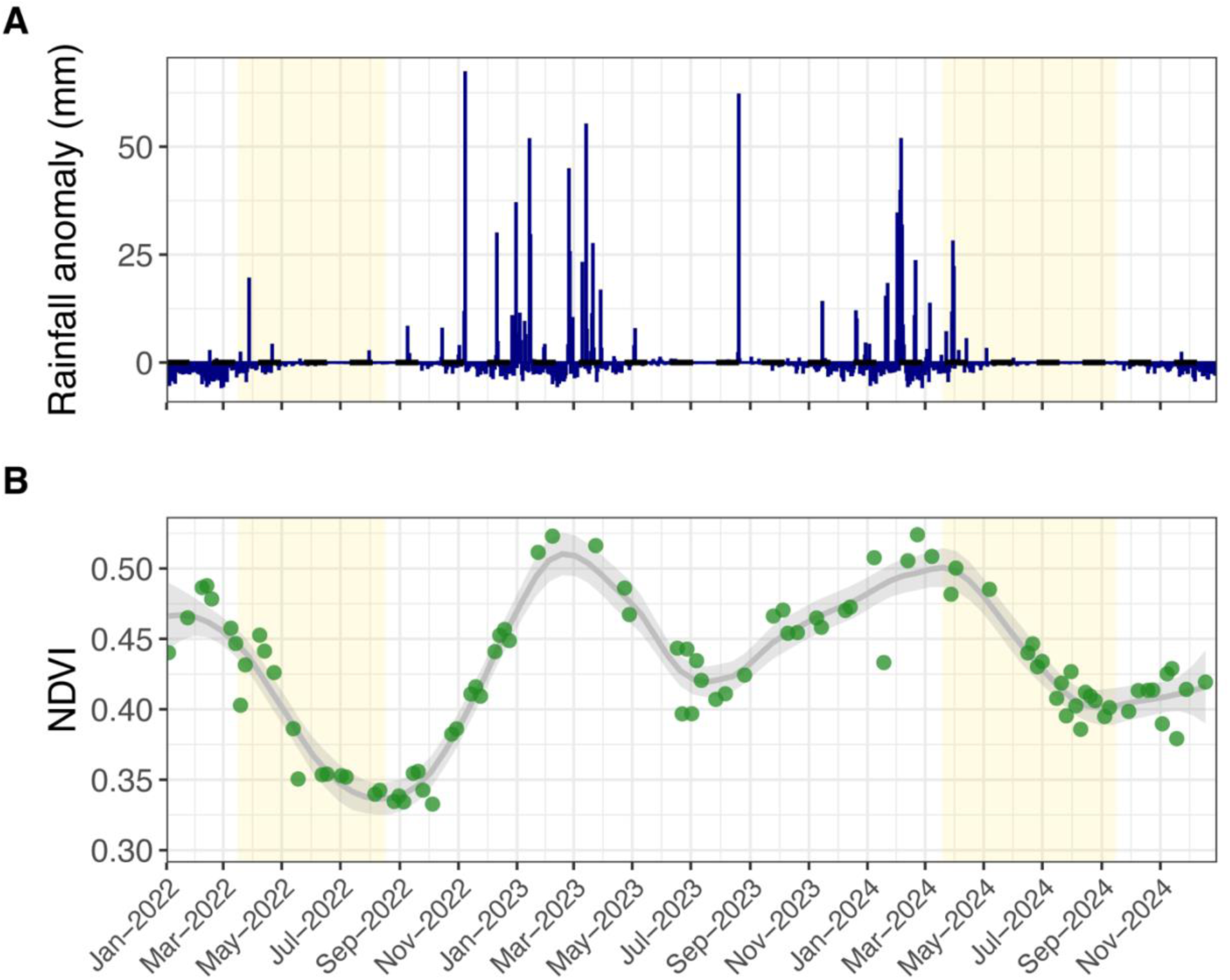
The “outbreak” of dysbiosis in 2024 followed a year with anomalously heavy winter precipitation, which likely increased the available floral resource supply for bumble bee populations. Yellow shaded areas show the sampling periods for *B. vosnesenskii* in 2022 and 2024. A) Atmospheric rivers resulted in high rainfall at the field site during the winter of 2022-2023. Shown is the historical mean (1981-2010) for each day of the year subtracted from the observed precipitation. Data were obtained from the PRISM Daily Spatial Climate Dataset. B) Plant productivity (and thus potential floral resources), as measured by Sentinel-2 normalized difference vegetation index (NDVI) data, was sustained at higher levels in 2023 versus 2022. NDVI data were obtained from an area centered on the field site with a radius equal to the expected foraging range of *B. vosnesenskii*. The gray line is a loess smoothing function.

Our proposed conceptual model to explain these results is that fluctuations in weather and floral resources in 2023 induced shifts in the *B. vosnesenskii* life cycle, in turn scrambling the seasonal trajectory of dysbiosis in 2024 (Fig. S5). Bumble bees are usually univoltine, with successive colony cycles separated by diapause and occurring in different nest sites^55^. However, their life history can be quite plastic in regions with mild winters and sufficient floral resources. Across various *Bombus* species, workers may persist and continue foraging throughout the winter; multiple colony cycles may occur without diapause; and old nests can be reused by newly produced queens^73,74,130,131^. For example, in a locality in the northwestern U.S. planted with a late-blooming crop, *B. vosnesenskii* appears to be bivoltine and capable of skipping diapause^132^. In coastal Southern California, which generally has warm and dry summers, life history plasticity may normally be constrained by limited floral resource availability.

We speculate that anomalously abundant resources during the summer of 2023 facilitated an extra fall colony cycle and/or overwintering of whole colonies (including workers). These life history responses would shorten or eliminate diapause and might circumvent nest replacement, allowing late-season dysbiosis to persist from the end of one year to the beginning of the next (Fig. S5). Such responses may be facultative, resulting in a composite bee population with different colony histories. This would help explain why interindividual microbiome variability was higher overall in 2024 versus 2022 (Fig. 2D) and why in 2024, males—which, in social insects, generally do not overwinter—began the season with low levels of environmental bacteria (Fig. 4A). However, direct observation of colonies in the field will be needed to evaluate this model.

Climate change is altering the timing and form of life cycles in many organisms^133,134^. In temperate-zone insects, unusually warm conditions can enable extra generations to be completed per year and increase overwintering survival^135,136^. These are the same kinds of weather-driven life history changes that, according to our conceptual model (Fig. S5), may have caused pervasive microbiome dysbiosis in the 2024 *B. vosnesenskii* population. Although only based on two years of sampling, the findings presented here tentatively illustrate the potential for extreme weather events—an increasingly common phenomenon with climate change—to regulate microbiome dynamics, potentially due to effects on host life cycles. These dynamics may also feed back to affect host life history plasticity. In the case of bumble bee dysbiosis, which is associated with disease, we speculate that it could constrain bumble bees’ ability to adjust their life history to optimally track changing environmental conditions.

## Conclusions

Our findings illustrate the challenge of interpreting microbiome dynamics in wildlife, particularly in the Anthropocene, as natural forces become difficult to disentangle from human influences. We show that dysbiosis is pervasive in a wild bumble bee community, including two at-risk species (*B. sonorus* and *B. californicus*), and argue that it is likely harmful to individual bee health. In other hosts, dysbiosis and related phenomena (e.g., coral bleaching) are often caused by anthropogenic stressors. In *B. vosnesenskii*, dysbiosis is not correlated with diet, foraging effort, or heat exposure. Further, it is notable that dysbiosis increased in prevalence following environmental conditions that were particularly conducive to bumble bee population growth. However, there are numerous other potential stressors that were not measured in this study, such as pesticides or viruses, and so the causal role of stress remains uncertain.

We propose that dysbiosis is widespread, in *B. vosnesenskii* as well as other *Bombus* species, because it is intrinsic to the annual life cycle of bumble bees. This idea is not at odds with dysbiosis being harmful to host health. Dysbiosis may evolve as a byproduct of tradeoffs between microbiome maintenance and other organismal functions, or due to relaxed selective pressure in post-reproductive and senescing colonies. And yet, even if dysbiosis is a typical feature of the host life cycle, it may not recur in the same way every generation. In the wild, host life cycles are not fixed but dependent on weather, resources, and other factors. We speculate that as the climate becomes more volatile, so too will life history, and in turn, microbiomes.

Testing these hypotheses will require both manipulative experiments—to which bumble bees are amenable—and longer-term microbiome data. Indeed, the strong seasonal and interannual variability we observe in the microbiome of *B. vosnesenskii* demonstrates how sampling in only one month or one year fails to capture “the microbiome” of a wild population. Intensive and sustained microbiome monitoring should be considered not just in bumble bees, but in any host with significant ecosystem impacts or conservation concern. Such data will be essential for teasing apart natural from anthropogenic microbiome dynamics, and more generally, for predicting and managing wildlife microbiomes.

## Materials & Methods

### Field collections

From March 16 to August 17, 2022, and March 19 to September 16, 2024, we collected *Bombus vosnesenskii* from the Niguel Botanical Preserve (NBP) in Laguna Niguel, Orange County, CA. NBP is a 7.3-hectare public garden with plants from Mediterranean climates, including those native to the range of *B. vosnesenskii* as well as non-native species. In 2024, we also collected *Bombus melanopygus*, *Bombus californicus*, and *Bombus sonorus*. Roughly every two weeks, several workers and males (when present) were collected while foraging with an insect net. We recorded the identity of the plant each bee was foraging on. Bees were transported to the lab in a cooler and anaesthetized on ice before removing the gut (midgut and hindgut) with ethanol-sterilized forceps. The rest of the body was stored at −20 °C. The gut was then homogenized with a sterile plastic pestle, suspended in 1 mL of 20% glycerol in phosphate-buffered saline, and stored at −20 °C. Forewings and one midleg were removed for trait analysis and genotyping, respectively.

### 16S rRNA gene sequencing & qPCR

To extract DNA from gut homogenates, we used the ZymoBIOMICS-96 kit, with the following modification to the lysis step. We bead-beat 250 µL of gut homogenate in 750 µL of Zymo Lysis solution with ∼0.5 mL autoclaved Zirconia silica beads (BioSpec). Bead beating was performed using a Benchmark Beadblaster 96 homogenizer at 2 minutes at 1800 rpm, rested for 1 minute, and 2 minutes at 1800 rpm. We included a total of three extraction blanks and three ZymoBIOMICS Microbial Community Standards. PCRs used barcoded universal primers targeting the V4 region of the 16S rRNA gene (515F/806R), following the V.1 protocol used by the Earth Microbiome Project^137^, with the only modifications being that reactions included 10.5 µL water and 12.5 µL Platinum HotStart PCR Master Mix 2X (Thermo Fisher Scientific), and that PCRs were performed in duplicate. One no-template control was included per PCR plate. Cleanup and normalization (up to 25 ng DNA per sample) were performed using the SequalPrep kit (Applied Biosystems). Pooled amplicons from 2022 samples were run on an Illumina Miseq (2 x 300 bp) at the UCI Genomics Research and Technology Hub. Amplicons from 2024 samples were run on an Illumina MiSeq (2 x 250 bp) at the CU-Boulder Center for Microbial Exploration Sequencing Center. Raw sequence data are available from NCBI (SRA BioProject #PRJNA1423801).

Processing of the 16S rRNA amplicon data followed a previously described protocol^43^. Initial steps were performed for each sequencing run (2022 versus 2024 samples) separately. From demultiplexed libraries, sequencing adapters and primers were removed with cutadapt^138^. Sequences were then trimmed and quality-filtered (maxEE = 2, truncQ = 2, 200 bp fwd, 150 bp rev), and denoised using DADA2^139^. The two ASV tables were then merged for further processing and analysis. Chimeras were removed and taxonomy was assigned to the merged ASV table using the RDP Naive Bayesian Classifier^140^ against the SILVA 138.2 SSU rRNA reference database^141^. The ASV table, metadata, and R code are available from Dryad (https://doi.org/10.5061/dryad.1jwstqkb8).

The ASV table, taxonomy, and metadata were imported into R and analyzed using mctoolsr (https://github.com/leffj/mctoolsr). Low-abundance ASVs (< 10 total reads across the dataset), and those without domain-level classifications were removed. Because 18S or mitochondrial 16S rRNA genes amplified by our primer pair can provide information about eukaryotic microbes^46^, we manually reclassified abundant nonbacterial ASVs using blastn searches against the NCBI nt database. This uncovered three ASVs matching with 100% sequence identity to 18S rRNA genes of well-known bumble bee parasites: the microsporidian *Nosema bombi* and the trypanosomatids *Crithidia expoeki* and *Crithidia bombi*. Using the same approach, we also manually reclassified a highly abundant bacterial ASV with no genus-level name as *Neokomagataea* based on a 100% identity match. Three abundant ASVs classified only to Enterobacteriales were also assigned genus names based on 99.2-100% identity matches.

For all analyses besides those concerning eukaryotic parasites, we filtered out all mitochondria, chloroplast, and eukaryotic sequences. We confirmed that the eight bacterial genera present in the ZymoBIOMICS Microbial Community Standard made up the eight most-common genera among our three mock community samples. Other taxa were present at lower abundances, likely reflecting cross-contamination from bee gut samples. Following established recommendations^142^, we used the tool decontam^143^ to identify putative reagent contaminants based on their prevalence among the six negative controls (DNA extraction blanks and no-template controls) from which we obtained sequence data. Gut samples with >10% contaminant sequences and negative control samples were removed. Contaminant ASVs were filtered out of the remaining samples. Finally, we randomly subsampled (i.e., rarefied) all samples to 1000 sequences. This step eliminated one *B. vosnesenskii* worker and one *B. melanopygus* worker from further 16S sequence analysis (except the parasite analyses, which used the pre-rarefaction dataset).

To infer the oxygen tolerance of the most prevalent bacterial genera in *B. vosnesenskii*, we used the BacDive database, a compilation of reported cultivation conditions for formally described bacterial isolates^144^. Data for two genera, *Schmidhempelia* and *Raoultella*, were absent from BacDive. *Schmidhempelia*, which has not been cultivated, was manually classified as an anaerobe based on metabolic properties inferred from its genome^145^. For *Raoultella*, we substituted values reported from its sister genus *Klebsiella*^146^.

We measured absolute abundance (16S rRNA gene copy number) using quantitative PCR on a qTOWER3 system (Analytik Jena) with universal 27F/355R primers, as described previously^43,147^. Copy numbers in qPCR reactions (with 1 μL template) were calculated from serial dilutions of plasmid DNA carrying the target gene and averaged per sample across three technical replicates. To estimate per-gut copy numbers, these values were multiplied by the volume of elution buffer used per DNA extraction (20 μL) and by 4, to account for the fact that extractions used ¼ of the total gut homogenate.

### Metagenomics

Based on the 16S rRNA gene amplicon data, we chose 8 core-dominated and 8 environmental-dominated *B. vosnesenskii* workers (all collected in 2022) for metagenomic sequencing. DNA was extracted from 250 uL of the gut homogenate described above, using the Zymo Quick-DNA Miniprep Plus extraction kit, with the “Fluids and Cells” protocol and overnight Proteinase K digestion. DNA was sent to the Joint Genome Institute (JGI) for library prep and sequencing on an Illumina NovaSeq with the S4 flow cell (2 x 200 bp reads). Sequence data are available from the JGI Genomes Online Database (GOLD Biosample IDs Gb0397770-Gb0397785).

JGI mapped the reads against a reference genome of *B. impatiens* (NCBI RefSeq: GCF_000188095.3) to remove host-derived reads. We ran the remaining reads through the SqueezeMeta pipeline v1.6.5, with default parameters for co-assembly^148^. The sequences were pooled and assembled with Megahit^149^, binned with CONCOCT^150^ and Metabat2^151^. Bin quality was assessed with CheckM^152^. We selected 8 medium or high-quality^153^ metagenome-assembled genomes (MAGs) with completeness 81.9-99.8% and contamination 0-2.0% for further analyses. Using the KBase platform^154^, we annotated MAGs with RASTtk v. 1.073^155^ and assigned taxonomy with GTDB-Tk v 2.3.2^156^. We used the PathoFact 2.0 pipeline to identify antimicrobial resistance genes, virulence factors, and toxins^157^.

### Host traits

To measure pollen-derived resources in the *B. vosnesenskii* gut, we counted the total number of pollen grains per gut using a hemacytometer under 100X magnification and two technical replicates per sample, following a previous protocol^40^. Body size variation in *B. vosnesenskii* was estimated using the length of the forewing marginal cell, an established proxy^109,119^. Left and right wing marginal cell lengths were recorded from wing scans taken on a Zeiss Stemi 508 microscope with an Axiocam 208 camera and averaged per individual. To infer lifetime foraging effort, we scored each *B. vosnesenskii* wing with an established, qualitative wing wear scale as 0 (fully intact), 1 (some minor indentations on the wing margin), 2 (indentations present on most of the wing margin), or 3 (> 5% of total wing area missing)^158^. Wing scores were summed per individual before analysis. Data are available in Dryad (https://doi.org/10.5061/dryad.1jwstqkb8).

### Host genotyping

To estimate sibship in *B. vosnesenskii* workers and males, we genotyped all individuals at eight established microsatellite loci (B124, BT10, B96, BT30, BTMS0066, BTMS0083, BTMS0059, and BMS0044), following previous work^159^. We extracted DNA by incubating macerated midleg tissue in 150 μL of 5% Chelex solution and 5 μL of Proteinase K for 1 h at 55 °C, 15 min at 99 °C, 1 min at 37 °C, and 15 min at 99 °C (Strange et al. 2009). Extracted DNA was PCR-amplified in two multiplexes (Plex 1: B124, BT10, B96, BT30; Plex 2: BTMS0066, BTMS0083, BTMS0059, BTMS0044). The final 10 μL reaction volume of each multiplex, containing 1 μL of template, 1× Promega (Madison, WI) reaction buffer, 0.6 mM dNTPs, 0.2 - 0.4 μM fluorescent 5’ dye-labeled primers, 1.4 mM MgCl_2_, 0.001 mg BSA, 0.4 units Taq polymerase (Promega, Madison, WI) and ddH_2_O to fill to volume. PCR cycling conditions were: 95 °C for 7 min, 30 cycles of 95 °C for 30 s, 53/55°C for 1 min 30 sec, 72 °C for 30 s, and 72°C for 10 min. PCRs were performed with and separated on an Applied Biosystems 3730xl automated sequencer (Thermo Fisher Scientific) along with a dye-labeled size standard (GeneScan 500 LIZ) at the Utah State University Center for Integrated Biosystems. Fragment length polymorphisms resulting from the sequencer were then scored using Geneious Prime 2021.0.1^160^.

Specimens with > 4 loci scored per individual were subjected to colony assignment using full-pedigree likelihood methods in COLONY v2.0^161^. Prior to colony assignment, ∼50% of the specimens collected in 2022 and 2024 were assessed for genotyping error by calculating mean error rate per locus (*e_l_*) as described previously^162^. Of the eight total loci we used across all specimens (B124, BT10, B96, BT30, BTMS0066, BTMS0083, BTMS0059, BTMS0044), all eight loci successfully amplified in the 2022 specimens, and six of the eight loci (B124, BT10, B96, BTMS0066, BTMS0083, BTMS0059) amplified in the 2024 specimens. Differences in the PCR amplification success of microsatellites may be due to template DNA variability or PCR error^163^. Accounting for genotyping error rates in the COLONY analysis allows for more robust estimates of colony assignment. We ran the COLONY analysis using full-likelihood method for haplodiploid species, assuming monogamy for both males and females. Specimens were considered full siblings if they shared ≥ 80% genotype similarity.

### Weather, NDVI, & phenology

We used the 4 km-resolution PRISM ANd dataset^164^ to collect precipitation and daily maximum temperature data for the Niguel Botanical Preserve for both a historical reference period (1981-2010) and the time frame spanning our field collections (2022-2024). Normalized Difference Vegetation Index (NDVI) data were obtained using the Harmonized Sentinel-2 Surface Reflectance image collection linked with the Sentinel-2 Cloud Score image collection^165^. Images with > 10% cloud cover were omitted, then pixels with a cloud score < 0.60 were filtered out. NDVI data at 10 m resolution were extracted and averaged from a buffer around the Niguel Botanical Preserve with a radius of 3.2 km, an estimate of the maximum foraging distance of *B. vosnesenskii*^76^. Both datasets were accessed via Google Earth Engine^166^.

The historically typical phenology of *Bombus vosnesenskii* in California was visualized using curated specimen records from 1880-2022 in the Bumble Bees of North America dataset (www.leifrichardson.org/bbna, accessed Oct. 12^th^ 2023). We included all records with the date of collection but regardless of caste (queen versus worker) or sex, because many of them lacked this information.

### Statistical analyses

All statistical analysis was conducted in R v. 4.5.0^167^. Several analyses (listed below) employed linear models (“lm” function in the stats package^167^) with the proportion of environmental bacterial sequences, sex, and the interaction as predictors. For models of alpha diversity, body size, and *Schmidhempelia* relative abundance, the interaction term was not significant, and the model was rerun without it to evaluate the significance of fixed effects. In models of pollen grain counts and 16S rRNA gene copy numbers, where the interaction term was significant, separate models were fitted for workers and males. In the latter models, the response variables were log-transformed to improve normality of residuals. In all cases, models were inspected to ensure that assumptions were met.

A variety of approaches were used in cases where linear models were not appropriate. To test whether oxygen tolerance differed between core and environmental bacterial genera, we first summarized data at the genus level by preserving the most common trait value among species within each genus. Three genera (*Gilliamella*, *Fructobacillus*, and *Arsenophonus*) had equal proportions of species belonging to two different trait values; these ties were resolved randomly to yield a single value per genus. Then, we used Fisher’s exact test (“fisher.test” function in the stats package^167^) on the resulting table (rows = core or environmental, columns = oxygen tolerance categories, values = number of genera) to test the null hypothesis of independence between rows and columns. To test whether dysbiosis predicts infection by parasites (*Crithidia bombi* and/or *Nosema bombi*), we converted parasite relative abundance into a binary variable (> 0 sequences = present) and then used a generalized linear model with a binomial error distribution (“glm” function in the stats package). To test whether dysbiosis predicts wing wear, an ordinal variable from 0-6, we fitted a cumulative link model using the “clm” function in the ordinal package^168^. For analyses of the effect of sex and collection date on the proportion of environmental bacteria, beta regression models were fitted using the “betareg” function in the betareg package^169^. Foraging patterns (in terms of plant families visited) were analyzed by training a random forest classifier (“randomForest” function in the randomForest package^170^) with 1000 trees and with the proportion of environmental bacteria and sex as predictors. Two individuals without recorded plant visitation data were excluded.

The vegan package^171^ was used for multivariate analyses. To visualize variation in microbiome composition among samples, we calculated Bray-Curtis dissimilarities for each year’s dataset (2022 versus 2024) and performed a non-metric multidimensional scaling ordination. These dissimilarities were also used to statistically test, with the “betadisper” function, whether within-group beta diversity (i.e., dispersion) varied as a function of microbiome type (core-versus environmental-dominated) and year. Tukey’s Honest Significant Difference method was used to test pairwise differences in mean dispersion. We used distance-based redundancy analysis, implemented with the “capscale” function, to test the effect of time and sex on microbiome composition (Bray-Curtis dissimilarities). Significance was assessed with a permutation test with 999 permutations.

## Acknowledgments

We thank the reviewers for comments that improved the manuscript, L. Richardson (Xerces Society for Invertebrate Conservation) for bumble bee records, S. Weber for advice on weather and NDVI analysis, T. Lindsay for assistance with microsatellite genotyping, L. F. Delgado for assistance with PathoFact, J. Schlauch for bee and flower icons, M. McDivitt and the Niguel Botanical Preserve for permission to conduct fieldwork, the UCI Research Cyberinfrastructure Center for computing support, and Willow Creek Farms for preliminary collections that inspired this study. B.S. acknowledges support from the National Science Foundation Postdoctoral Research Fellowships in Biology Program (#2305844). Metagenomic sequencing was provided by the Joint Genome Institute’s Community Science Program (#509743). This material is based upon work supported by the National Science Foundation (#DEB-2443951).

## Supplemental Material

**Table S1.** Metagenome-assembled genomes (MAGs) from the gut of wild *B. vosnesenskii* workers collected in 2022. MAGs were classified to genus using the Genome Taxonomy Database (GTDB) Toolkit. The isolation source of the top hit (by average nucleotide identity) in GTDB is listed. “Category” indicates whether a given bacterial genus is categorized as core or environmental for microbiome analyses.

| MAG ID | MAG genus | Source of closest match in GTDB | Category |
| --- | --- | --- | --- |
| BIF_1 | *Bifidobacterium* | *Bombus* spp. (adult gut) | Core |
| BIF_2 | *Bifidobacterium* | *Bombus* spp. (adult gut) | Core |
| BOM_1 | *Bombiscardovia* | *Bombus* spp. (adult gut) | Core |
| SNO_1 | *Snodgrassella* | *Bombus* spp. (adult gut) | Core |
| SCH_1 | *Ca. Schmidhempelia* | *Bombus* spp. (adult gut) | Core |
| GIL_1 | *Gilliamella* | *Bombus* spp. (adult gut) | Core |
| ARS_1 | *Arsenophonus* | *Apis mellifera* (adult hemolymph) | Environmental |
| FRU_1 | *Fructobacillus* | *Apis mellifera* (diseased larva) | Environmental |

**Figure S1.**
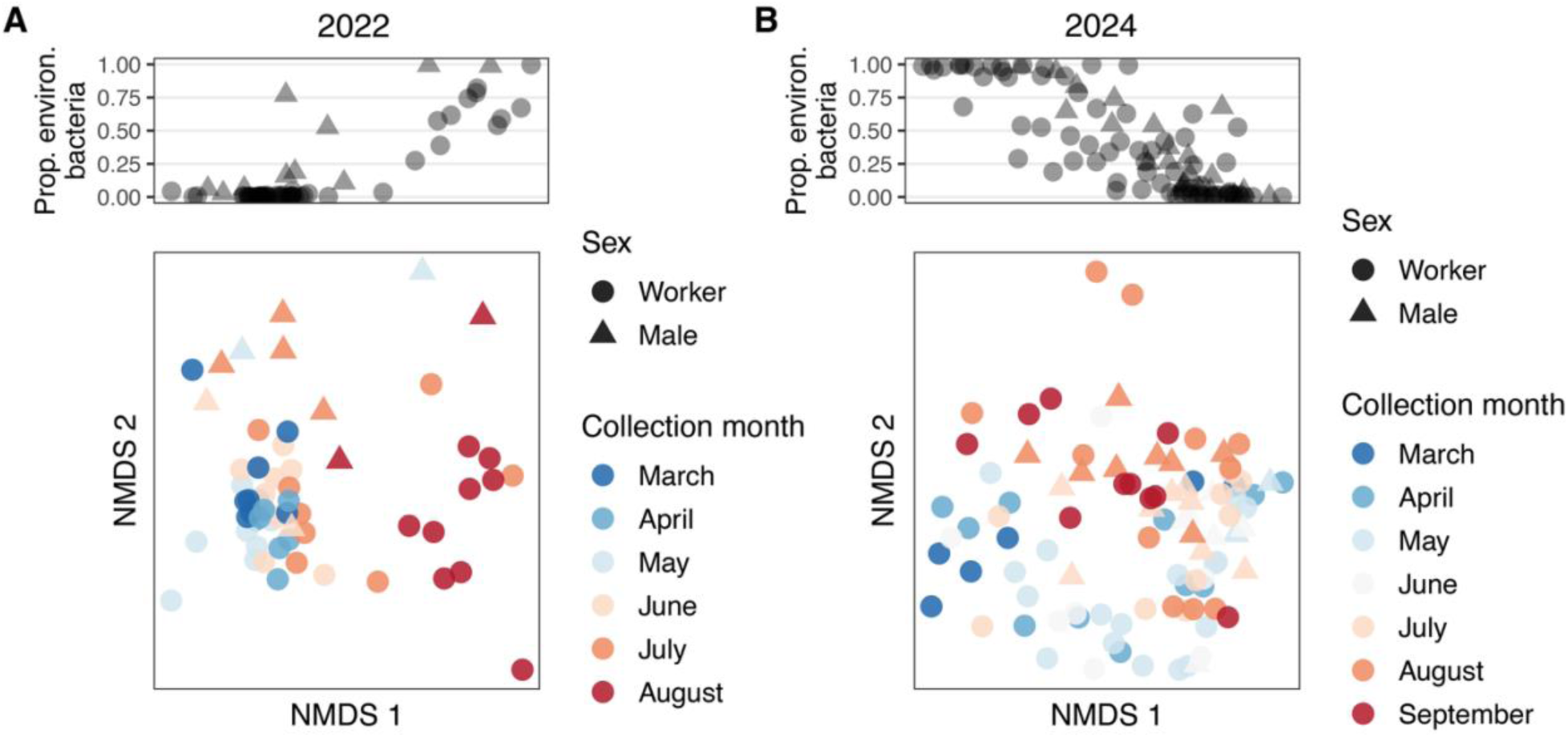
*B. vosnesenskii* gut microbiome composition varies over the season and between workers and males. A) Nonmetric multidimensional scaling (NMDS) ordinations of Bray-Curtis dissimilarities in the 2022 sample set. The upper panel shows the association between a sample’s position on the first NMDS axis and its proportion of sequences belonging to environmental bacteria. B) The same as panel A, for bees sampled in 2024.

**Figure S2.**
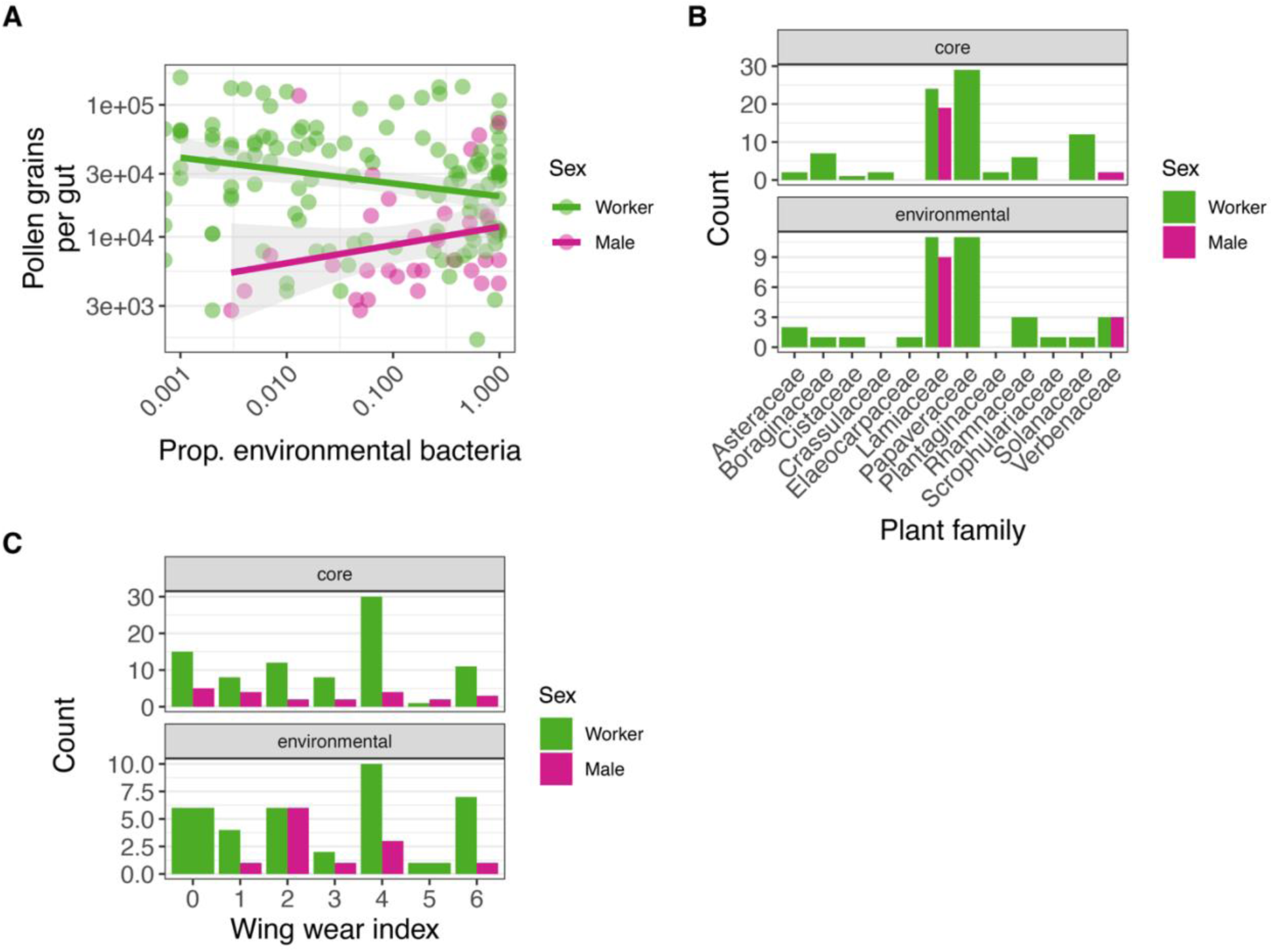
Three potential stressors do not predict dysbiosis in *B. vosnesenskii*. Samples from both 2022 and 2024 are shown together. A) Pollen resources were less abundant in the gut of males versus workers, but were not significantly associated with the relative abundance of environmental bacteria. To quantify pollen in gut contents, exines (which are indigestible) were counted using microscopy. Lines show linear model fits and 95% confidence intervals. B) Bees with dysbiotic microbiomes did not collect resources from an obviously altered or restricted set of plant taxa. Shown are the plant families on which bees were foraging when they were collected. To facilitate visualization, bees were binned based on microbiome composition into majority-core or majority-environmental groups. C) Dysbiosis was not associated with intensive foraging. An established wing wear scoring system was used to infer past foraging effort. The index shown for a given individual is the sum of left and right wing scores; odd scores are less common because wings tended to have similar wear. Bees were binned for visualization as in panel B.

**Figure S3.**
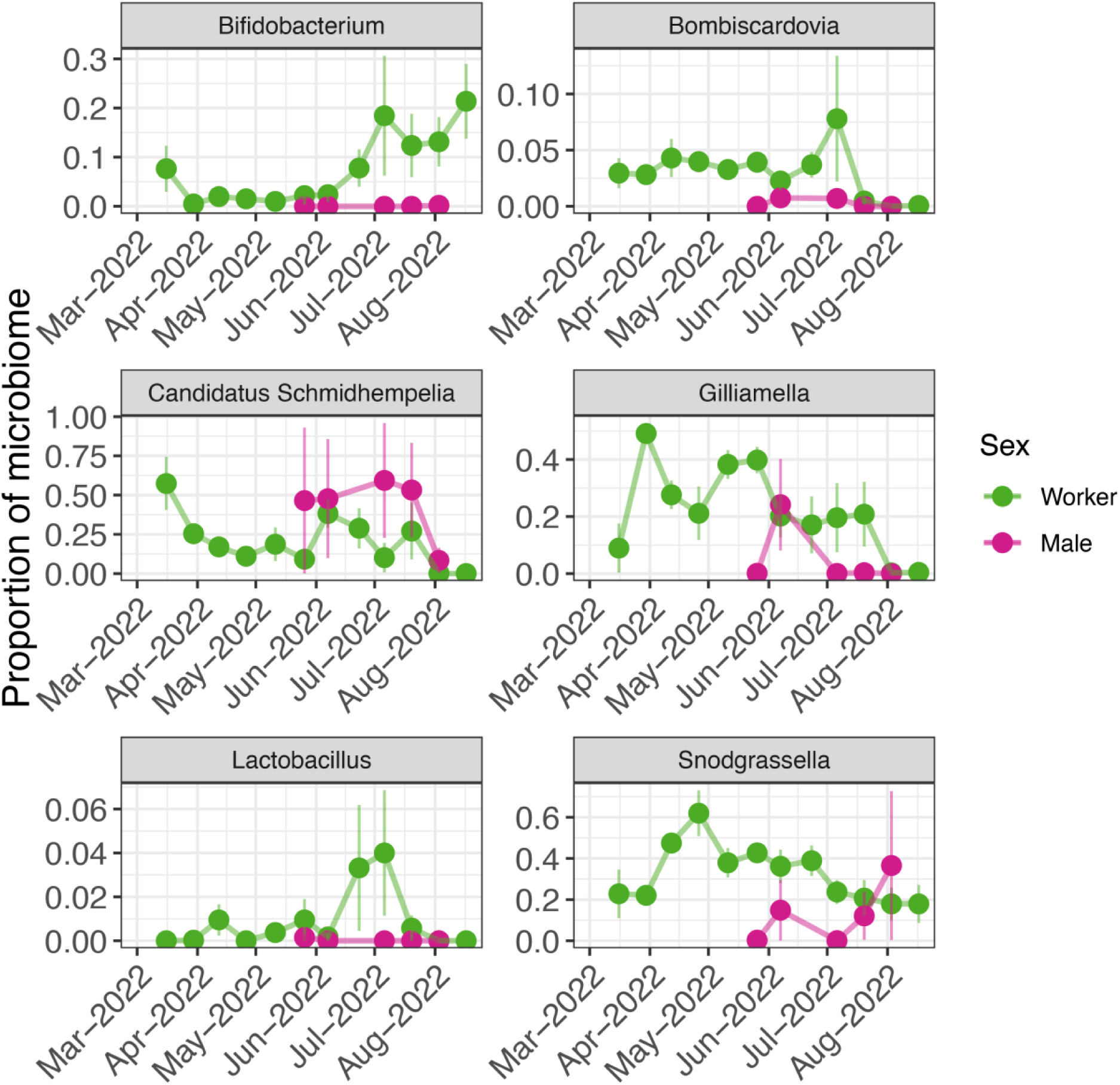
Relative abundances of individual core (bumble bee-specialized) gut bacterial taxa in *B. vosnesenskii* collected in 2022 vary seasonally and between taxa. The y axis shows the proportion of each taxon out of the total bacterial community. Points represent means of replicate bees sampled on the same day and vertical lines represent the SEM (N = 50 workers, 10 males). Tick marks on the x axis correspond to the first day of each month. Note that the y axis scale differs among taxa. One core genus, *Bombilactobacillus*, is not shown because it was absent from *B. vosnesenskii* collected in 2022.

**Figure S4.**
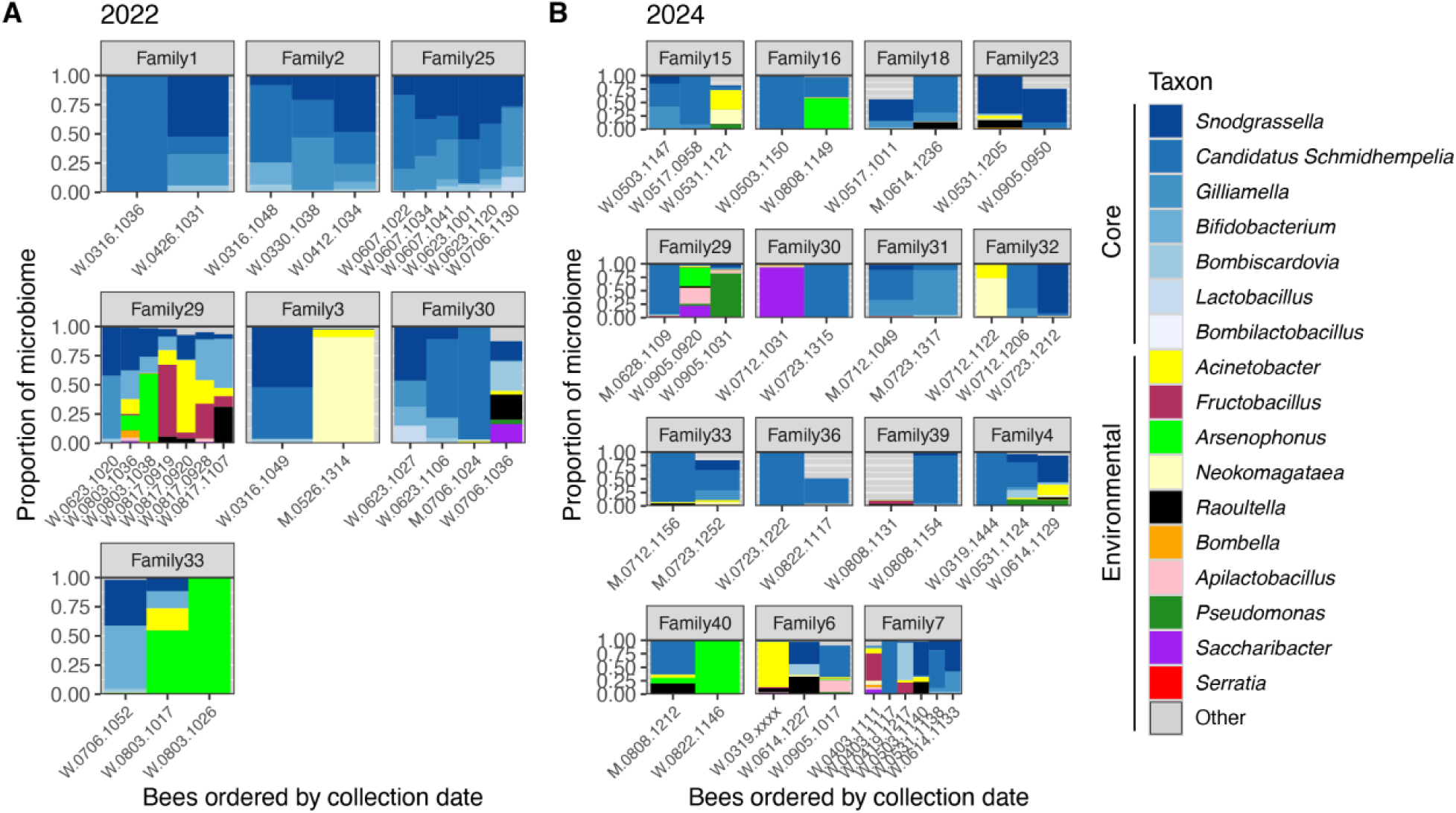
Evidence that the microbiome of *B. vosnesenskii* can vary within colonies over time (i.e., between nestmates sampled at different time points). Siblings were inferred from microsatellite genotyping. Family groups (i.e., colonies) were reconstructed only within each year’s dataset and hence do not overlap between years, even if they share the same name. Only family groups with 2+ individuals are shown. Each column is an individual bee, labeled by sex (W = worker, M = male) followed by collection date (MMDD) and time of day. “Other” (gray) represents all environmental bacterial genera with <1% mean relative abundance across all samples. A) *B. vosnesenskii* sampled in 2022. B) *B. vosnesenskii* sampled in 2024.

**Figure S5.**
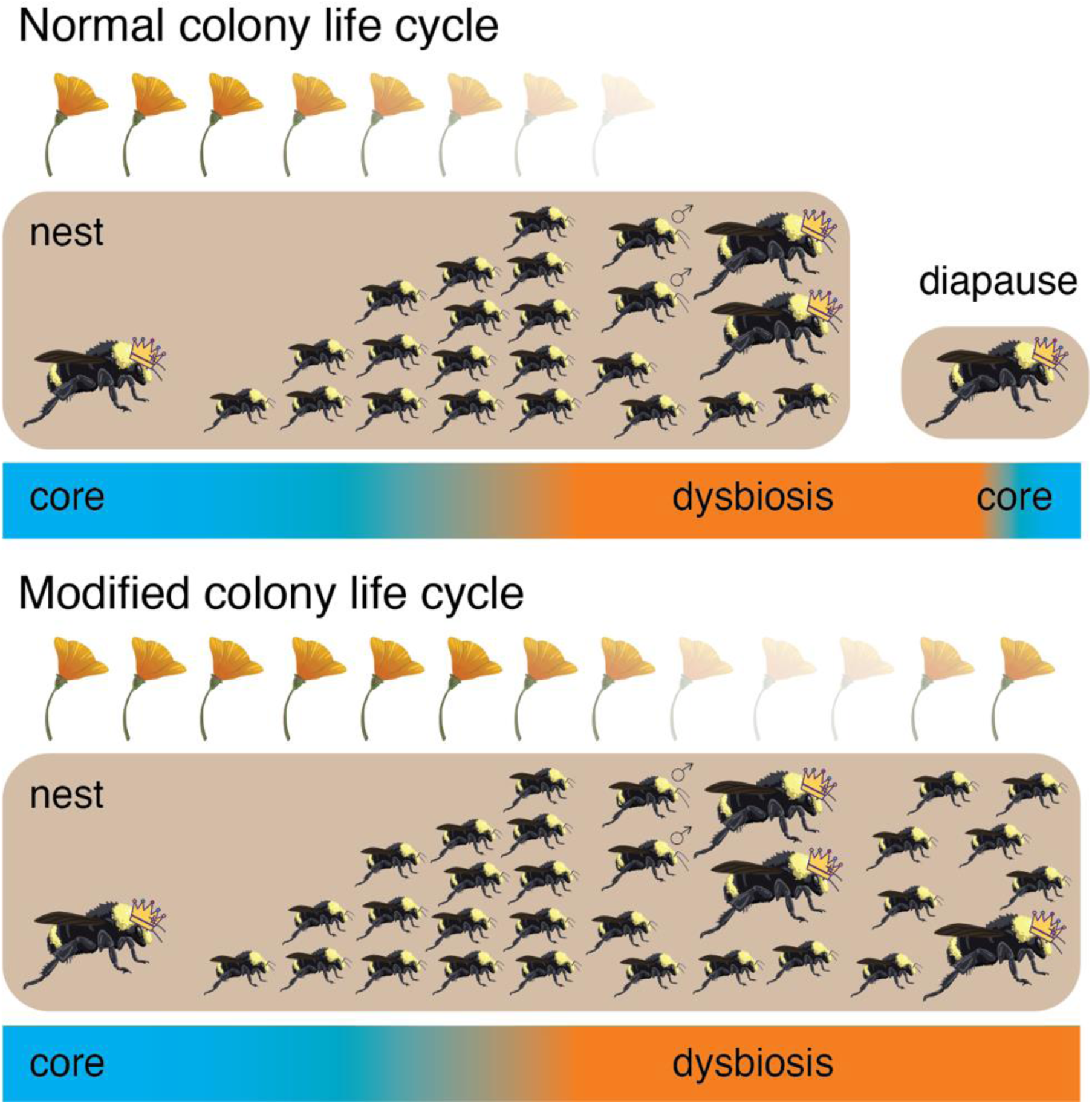
A proposed conceptual model to explain seasonal and interannual dynamics of the bumble bee gut microbiome. Top: a typical seasonal progression from core-dominated to dysbiotic microbiomes. In the standard annual *Bombus* life cycle, colony growth is followed by production of males and new queens, which mate; workers and males die, and the new queens undergo diapause and found new nests in the spring. Diapause and nest replacement reset the microbiome from year to year. We suggest that 2022 was representative of a relatively typical year for the focal *B. vosnesenskii* population, because similar temporal microbiome patterns have been reported in other locations and species. Bottom: sustained floral resources allow old nests (potentially even including workers) and thus dysbiosis to persist into the following year. We suggest that this dynamic occurred in 2023, resulting in widespread dysbiosis in 2024. Other types of life history plasticity have been documented in bumble bees (e.g., multivoltinism) and may have also occurred in the *B. vosnesenskii* population (not shown).

